# Tau protein aggregation induces cellular senescence in the brain

**DOI:** 10.1101/369074

**Authors:** Nicolas Musi, Joseph M. Valentine, Kathryn R. Sickora, Eric Baeuerle, Cody S. Thompson, Ashley Zapata, Qiang Shen, Miranda E. Orr

## Abstract

Tau protein accumulation is the most common pathology among degenerative brain diseases, including Alzheimer’s disease (AD), progressive supranuclear palsy (PSP), traumatic brain injury (TBI) and over twenty others^1^. Tau-containing neurofibrillary tangle (NFT) accumulation is the closest correlate with cognitive decline and cell loss, yet the mechanisms mediating tau toxicity are poorly understood. NFT-containing neurons do not die, which suggests secondary mechanisms are driving toxicity^2^. We evaluated gene expression patterns of NFT-containing neurons microdissected from AD patient brains^3^ and found they develop an expression profile consistent with cellular senescence described in dividing cells. This complex stress response induces a near permanent cell cycle arrest, adaptations to maintain survival, cellular remodeling, and metabolic dysfunction^4^. Moreover, senescent cells induce chronic degeneration of surrounding tissue through the secretion of pro-inflammatory, pro-apoptotic molecules termed the senescence-associated secretory phenotype (SASP)^5^. Using transgenic mouse models of tau-associated pathogenesis we found that NFTs induced a senescence-like phenotype including DNA damage, karyomegaly, mitochondrial dysfunction and SASP. *Cdkn2a* transcript level, a hallmark measure of senescence, directly correlated with brain atrophy and NFT load. This relationship extended to postmortem brain tissue from humans with PSP to indicate a phenomenon common to tau toxicity. Tau transgenic mice with late stage pathology were treated with senolytics to remove senescent cells. Despite the advanced age and disease progression, senolytic treatment reduced total NFT burden, neuron loss and ventricular enlargement; and normalized cerebral blood flow to that of non-transgenic control mice. Collectively, these findings indicate that NFTs induce cellular senescence in the brain, which contributes to neurodegeneration and brain dysfunction. Moreover, given the prevalence of tau protein deposition among neurodegenerative diseases, these findings have broad implications for understanding, and potentially treating, dozens of brain diseases.

---

In human AD, tau-containing NFT density closely tracks with disease severity^2^. However, NFTs do not induce immediate cell death^6^; instead *in silico* modeling predicts that NFT-containing neurons may survive decades^7^. To gain insight into how NFT-containing neurons endure an environment that is toxic to histologically adjacent cells^2,6^, we queried the publicly available GEO Profiles database^8^ for gene sets specific to NFTs. We evaluated laser capture microdissected cortical neurons containing NFTs from AD brains (GEO accession GDS2795) and compared them to adjacent histopathologically normal neurons for a within-subjects study design^3^. NFT containing neurons upregulated genes involved in cell survival and viability, inflammation, cell cycle progression and molecular transport and downregulated apoptosis, necrosis and cell death pathways (Figure 1a). NFκB, a pro-survival master transcriptional regulator of inflammation, was the highest predicted upstream regulator of the NFT-gene expression profile. In agreement with inflammatory activation, other predicted upstream regulators included IFNG, TNF, TLR4, IL1B and CXCL1 (Figure 1b). Collectively, the molecular pathways identified in the NFT analyses resembled cellular senescence. This complex stress response is detrimental to the local environment through the secretion of toxic, pro-apoptotic, inflammatory molecules (SASP); simultaneously senescent cells acquire a pro-survival gene expression pattern to protect themselves against their own toxic factors^5^.

**Figure 1.**
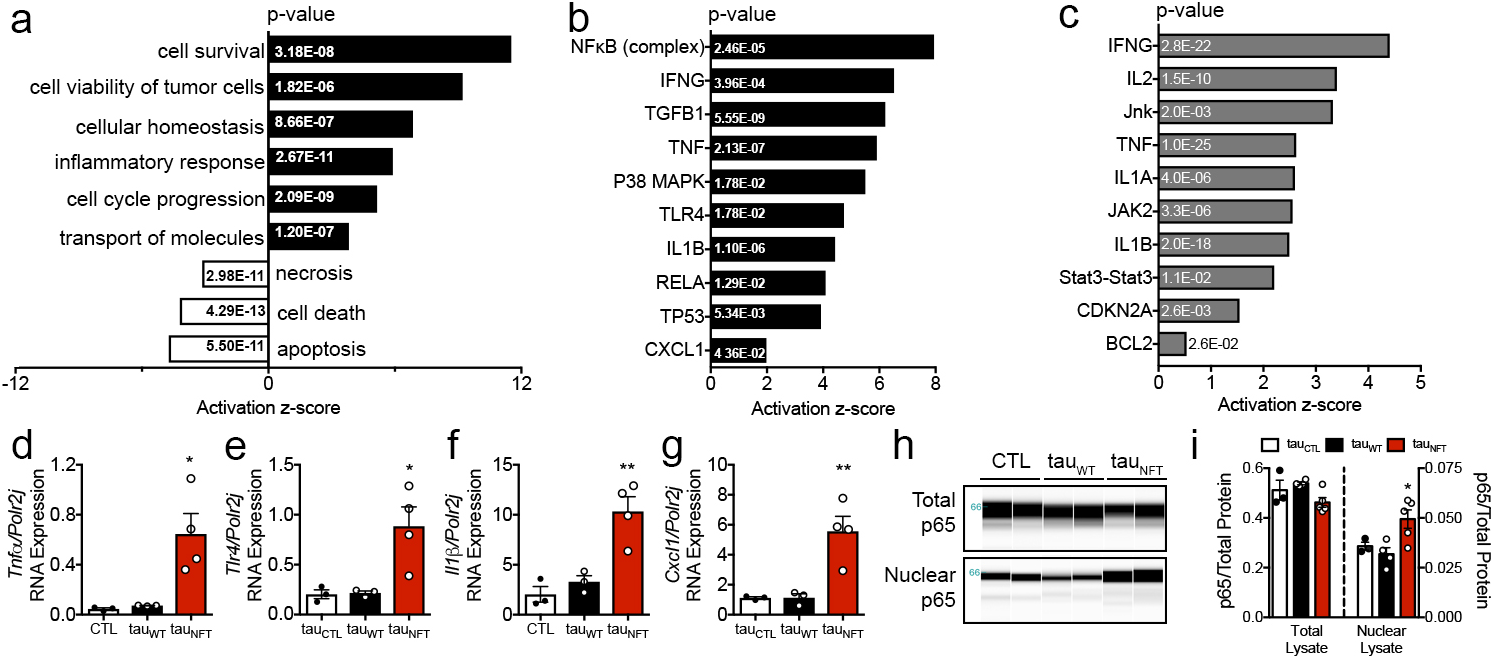
Neurofibrillary tangles upregulated cellular senescence-associated gene pathways in human Alzheimer’s disease neurons and tau transgenic mouse brains. (a) Pathways and predicted upstream regulators identified by Ingenuity Pathway Analyses (IPA, QIAGEN) as significantly enriched in Alzheimer’s disease patient-derived neurons with neurofibrillary tangles compared to non-tangle containing neurons; z-score plotted on x-axis and (p-value) indicated in bar graph. Cellular functions and (b) predicted upstream regulators employed by neurofibrillary tangle containing neurons derived from Alzheimer’s disease patient are shown. (c) Predicted upstream regulators of gene transcription in tau_NFT_ mice after the onset of neurofibrillary tangles (~6-mo-old vs. ~2-mo-old); z-score plotted on x-axis and (p-value) indicated in bar graph. (d-g) Quantitative gene expression on RNA isolated from CTL (open bar, n=3), tau_WT_ (closed bar, n=3) and tau_NFT_ (red bar, n=4) mouse forebrain targeting senescence-associated genes in pathways discovered in panels a-c; (d): *Tnfa*, *P* = 0.0114; (e) *Tlr4*, *P* = 0.0144; (f) *Il1b, P* = 0.0025; and (g) *Cxcl1, P* = 0.0040. (h) Immunoblot generated by capillary electrophoresis on subcellular fractionated mouse forebrain homogenate probed with anti-NFκB p65 antibody. Total cellular p65 (top blot) and nuclear localized p65 protein levels (bottom blot) were (i) normalized to total protein content. Total p65; *P* = 0.0758; nuclear p65, *P* = 0.0223. CTL: open bar, n=3; tau_WT_: closed bar, n=4; tau_NFT_: red bar, n=5. In all experiments, mice were aged 16-18-months-old; both males and females were included. Data are graphically represented as mean ± s.e.m. Significance determined by one-way ANOVA Tukey’s multiple comparison *posthoc*.

To investigate a link between NFTs and a senescence-like phenomenon in neurodegeneration, we used the rTg(tau_P301L_)4510 transgenic mouse line, hereon referred to as “tau_NFT_.” They develop well-characterized, aggressive, tau pathology in forebrain regions concomitant with neurodegeneration and cognitive deficits^9^ (pathology illustrated in Extended Data Figure 1). Mice that overexpress wild type human tau, “tau_WT_,” acquire age-dependent tau pathogenesis at a much slower rate, and are used to identify effects of elevated pre-pathogenic tau^10^; age-matched tau_NFT_ littermate mice without human tau overexpression serve as wild type controls, “CTL”.

To determine if NFT containing neurons in mice induced a gene expression profile resembling cellular senescence as seen in our human AD data, we assessed hippocampal gene expression patterns in tau_NFT_ mice before (~2-month-old) and after (~6-month-old) NFT formation (GSE56772). Consistent with NFTs from human AD, mouse NFTs also caused significant activation scores for IFNG, TNF, IL-1B, as well as enrichment in other senescence associated JAK, STAT, CDKN2A and BCL2 predicted upstream regulators (Figure 1c). Senescent cell numbers increase with age in regenerative tissues and contribute to functional decline and decreased lifespan^11,12^. Likewise, advancing age is the greatest risk factor for developing AD. To test the contribution of aging on cellular senescence in tau transgenic mice, we evaluated hallmarks of senescence in older mice (16-month-old tau_NFT_ and control mice). We found significantly elevated expression of the same SASP-associated genes in 16-month-old tau_NFT_ brains as identified in human AD NFT neurons, i.e., in 16-month-old tau_NFT_ brains, *Tnfa* was 13-and 8-fold higher than CTL and tau_WT_, respectively; *Tlr4* was 3-fold higher than both control genotypes; *Il1b* was 4-and 2-fold higher than CTL and tau_WT_, respectively; and *Cxcl1* was 4-fold higher than both control genotypes (Figure 1d-g). NFκB regulates the pro-survival, pro-inflammatory SASP gene expression profile characteristic of cellular senescence^13^. Consistent with NFκB pathway activation, the SASP profile and our transcriptomic data from mice and humans with NFTs, nuclear localized NFκB p65 was significantly increased in tau_NFT_ brains (Figure 1h-i).

Mitochondrial dysfunction is obligatory for SASP production and cellular senescence^14,15^. To examine mitochondrial bioenergetics, we performed high-resolution respirometry to yield accurate quantitative measurements of oxidative phosphorylation in response to specific substrates for complex I, complex II, fat oxidation and electron-transfer system (ETS) capacity. Across genotypes, we compared cortex, hippocampus and cerebellum. This allowed for evaluation of specific differences in oxygen consumption due to elevated transgenic tau (comparing CTL with tau_wt_ and tau_NFT_), pathogenic tau specific effects (comparing tau_wt_ to tau_NFT_), as well as the interaction among brains regions and tau expression (e.g., cortex and hippocampus express transgenic tau and develop NFTs, but cerebellum does not). We found a significant genotype main effect for oxygen flux in both cortex and hippocampus, indicating that global respiratory capacity was impaired in NFT containing brain regions (*P* < 0.0001; Figure 2), an effect primarily driven by CI+CII respiration coupled to ATP production (cortex: *P* = 0.0034; hippocampus: *P* = 0.0215; Figure 2g and h, respectively), and uncoupled or maximum respiratory capacity (cortex: *P* = 0.0248; hippocampus: *P* = 0.0261; Figure 2g and h, respectively). These changes were different between tau_NFT_ and each of the control mouse lines, CTL and tau_WT_ mice. Because tau_WT_ and tau_NFT_ mice express comparable total tau levels, alterations to respiratory capacity cannot be attributed to tau overexpression. Citrate synthase activity is a surrogate marker of total mitochondrial content/mass, and was similar across genotypes and brain regions (Extended Data Figure 2a) suggesting that the defects in cellular respiration were due to altered mitochondrial quality, not content/mass. Moreover, tau_NFT_ cerebellum did not show deficits in cellular respiration (Extended Data Figure 2b), indicating that mitochondrial dysfunction was present only in brain regions with persistent pathogenic tau expression.

**Figure 2.**
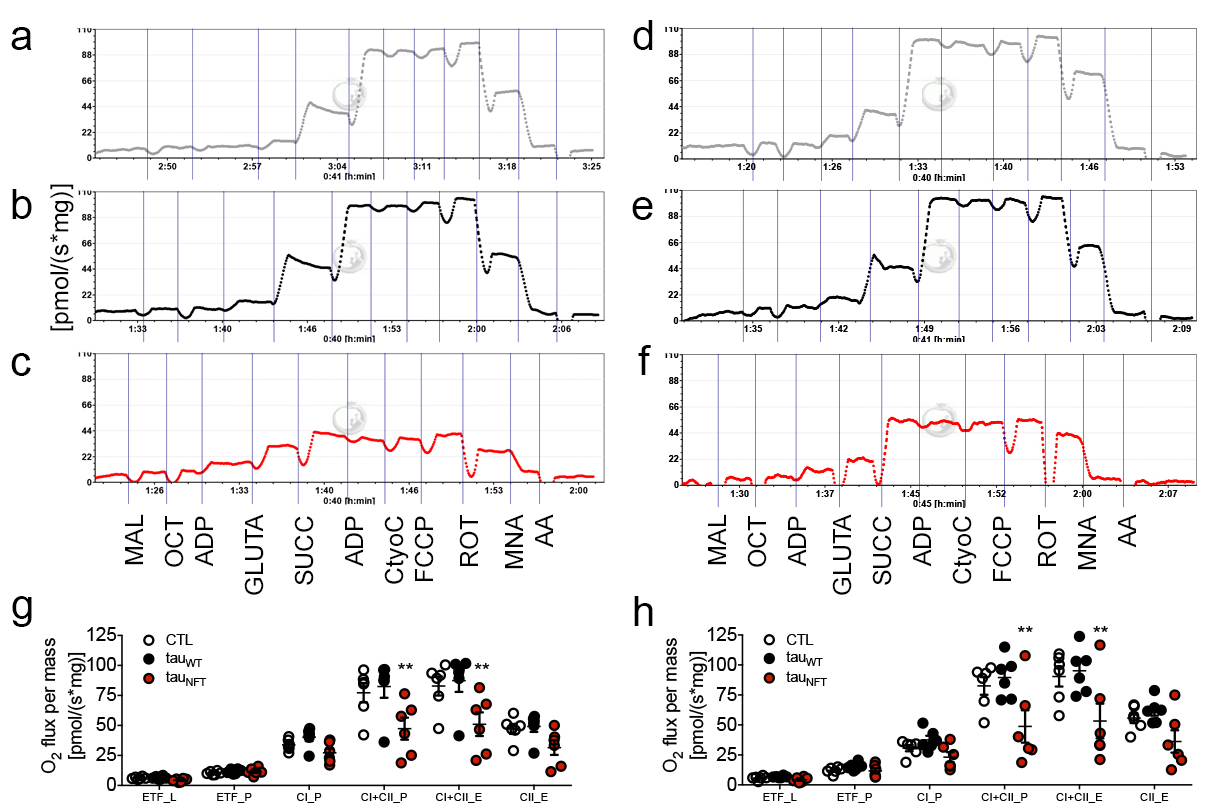
Neurofibrillary tangle-containing brain regions had impaired cellular respiration. (a-c) Representative respirometric traces from cortical and (d-f) hippocampal tissues illustrate genotype-specific differences in oxygen consumption using the SUIT protocol (top gray traces: CTL; black middle traces: tau_WT_; bottom red traces: tau_NFT_). (g) Tissue mass-specific respiration analyses indicate significant deficits in cortical and (h) hippocampal ATP-dependent and maximum oxygen consumption in mice with neurofibrillary tangle pathology. (ETF_L (fat oxidation in absence of ADP (state 2)), ETF_P (fat oxidation coupled to ATP production), CI_P (complex I activity linked to ATP production (state 3)), CI+CII_P (complex I & II linked respiration (state 3)), CI+CII_E (complex I & II linked respiration uncoupled (maximum respiration), CII_E (complex II activity uncoupled). Experimental mice were aged 16-18-months-old with n=6/group; both males and females were included. Data are graphically represented as mean ± s.e.m. Two-way ANOVA Tukey’s multiple comparison *posthoc*: ** *P* < 0.005.

Senescence-inducing stressors often inflict DNA-damage that occurs concomitant with changes in cellular function and morphology. Tau_NFT_ mouse brains displayed significantly elevated histone γ-H2ax, a highly sensitive marker of both double-stranded DNA breaks and cellular senescence^16^ (*P* = 0.0056; Figure 3a-b). Morphological and functional changes accompanying DNA damage may include increased cell size; altered nuclei that are either enlarged (karyomegaly) or syncytia (multinucleated)^4^; and/or altered lysosomal hydrolase activity, termed senescence associated (SA) beta-galactosidase (SA β-gal; activity at pH 6.0)^17^. We previously reported that neuronal stem-cell containing cultures derived from tau_NFT_ mice developed a “senescent-like” enlarged morphology *in vitro*^18^. Here we assessed the senescence-associated karyomegaly phenotype by analyzing nuclei area from NFT-containing cortical neurons, neighboring neurons without NFTs, and CTL cortical neuronal nuclei. NFT-containing neuronal nuclei were larger than non-NFT containing neurons in histologically adjacent tissue (8.5%, *P* = 0.0004) and CTL nuclei (7.9%, *P* = 0.0001) (Figure 3c-e). We repeated this experiment using histone 3, instead of DAPI, as a marker of nuclear size; tau_NFT_ nuclei were reproducibly larger than non-NFT containing neurons (*P* < 0.0001; Extended Data Figure 3) indicating that NFTs induce nuclear enlargement.

**Figure 3.**
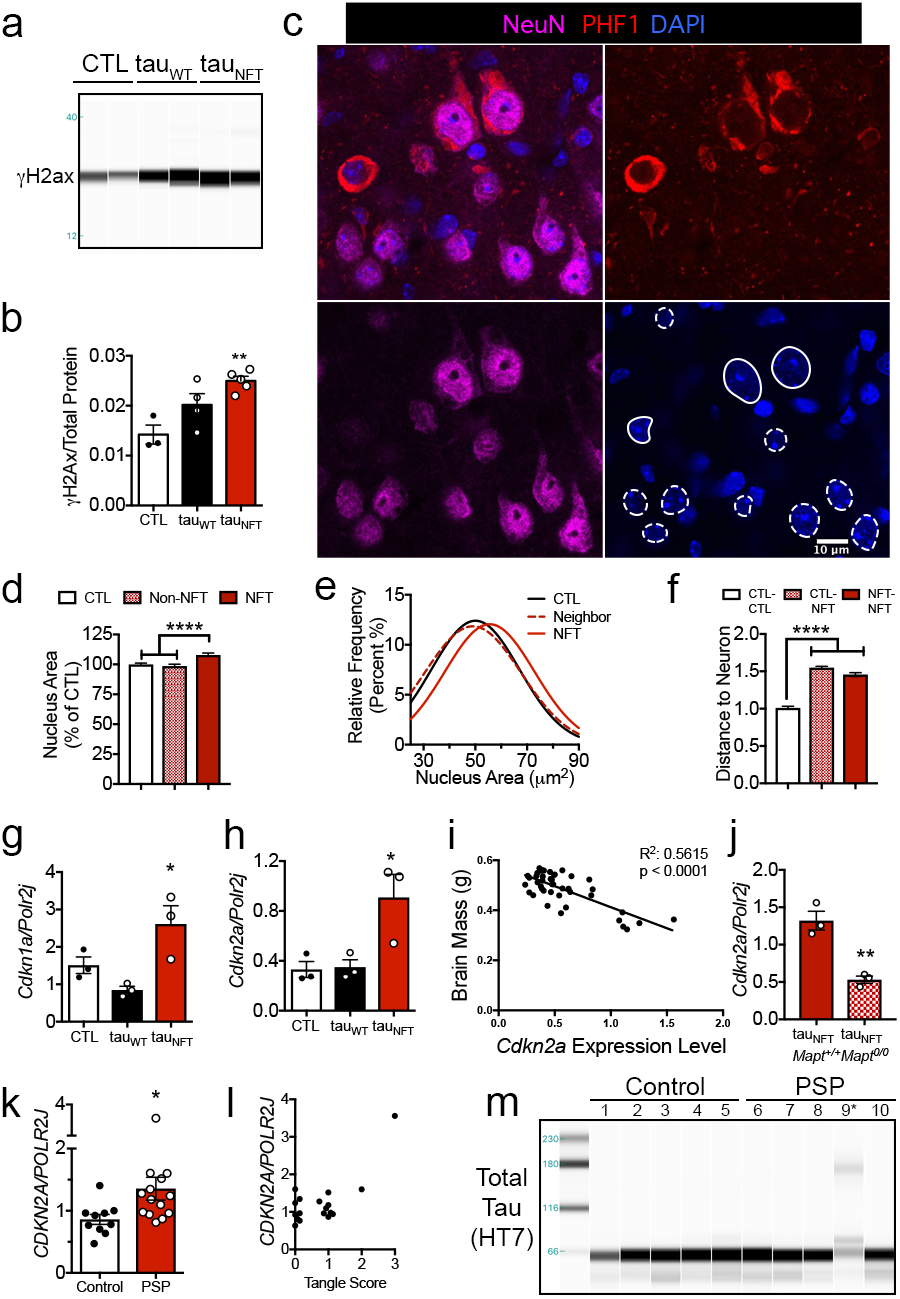
Neurofibrillary tangles induced senescence-associated DNA damage, neuronal karyomegaly, toxicity, and cell cycle dysfunction in mice and human patients. (a) Representative immunoblot generated by capillary electrophoresis on chromatin-bound fractions from mouse forebrain homogenate probed with anti-γ-H2ax antibody. (b) Densitometric normalization of γ-H2ax to total protein content (CTL: n=3; tau_WT_ n=4; tau_NFT_: n=5; ANOVA, *P* = 0.0056). (c) Immunofluorescence of cortical tissue using anti-NeuN to identify neurons (magenta), anti-PHF1 to identify neurofibrillary tangles (red) and DAPI to identify cell nuclei (blue). (d) Quantification of nuclear DAPI area from neurons without pathology or with neurofibrillary tangles (open arrow and red broken circle in Panel a from n = 4 CTL mice (total neurons counted = 641) and n = 3 tau_NFT_ mice (total neurons counted without NFTs: n = 346 and with NFTs: n = 346). ANOVA Tukey’s *posthoc*: CTL vs. NFT, *P* = 0.0001; NFT vs. nearest non-NFT neuron (non-NFT), *P* = 0.0004. (e) A Gaussian distribution derived from the frequency of cells (y-axis) plotted against the nuclei area in μm^2^ (14 frequency bins generated for each neuronal nuclei category n = 346 NFT neurons, n = 346 non-NFT neighbor neurons counted; n = 621 control neurons). Scale bar = 10μm. (f) We measured the distance between control neurons without neurofibrillary tangles (CTL, outlined with dotted white circles in panel c), CTL neurons to neurofibrillary containing neurons (NFT, outlined with solid white circles in panel a) and between two NFTs. Total counts: n = 1098 CTL to CTL neurons; n = 3809 CTL to NFT neurons; n = 1541 NFT to NFT neurons from n = 3 tau_NFT_ mice. ANOVA, **** *P* < 0.0001. (g) Quantitative gene expression analyses on RNA isolated from CTL (open bar, n=3), tau_WT_ (closed bar, n=3) and tau_NFT_ (red bar, n=3) mouse forebrain for *Cdkn1a, P* = 0.0207, and (h) *Cdkn2a* expression, *P* = 0.0216. (i) *Cdkn2a* expression levels were significantly correlated with brain atrophy (R^2^ = 0.5615, *P* < 0.0001; n = 43). (j) *Cdk2na* gene expression comparison between age-matched mice carrying tau_NFT_ on a *Mapt* wild type or *Mapt* knockout background. Unpaired two-tailed t-test, *P =* 0.0041; n=3/group). (k) *CDKN2A* gene expression of RNA extracted from post-mortem brain tissue from control older adult humans (n=10) was significantly lower than that of age-matched progressive supranuclear palsy (n=14). Unpaired two-tailed t-test, *P* = 0.0415. (l) *CDKN2A* expression plotted against neurofibrillary tangle deposition in the parietal lobe (ANOVA, *P* = 0.0008; Kendall’s Tau rank correlation *P =* 0.059). (m) Representative immunoblot generated by capillary electrophoresis on cortical brain homogenate from control and progressive supranuclear palsy human brains probed with total tau antibody, HT7. The individual with the highest *CDKN2A* expression (panel g) displayed high molecular weight tau, lane 9*. Data are graphically represented mean ± s.e.m.

In regenerative tissues and *in vitro* cultures, senescent cells may exhibit SA β-gal activity^19^; however, the consequence of SA β-gal activity in post-mitotic neurons is less clear. For example, SA β-gal activity has been reported in hippocampal neurons of young mice^20^. Similarly, SA β-gal reactive neurons were detected in one-month-old mice here (Extended Data Figure 4a, b). Examination of the hydrolase coding gene, galactosidase beta (β) 1 (*Glb1*), (Extended Data Figure 4c) revealed that tau_NFT_ mice expressed higher *Glb1* gene expression than controls, yet fewer tau_NFT_ hippocampal neurons displayed β-gal hydrolase activity at pH 6.0. Further, the number of SA β-gal reactive cells was positively correlated with brain mass (R^2^: 0.4852, *P* = 0.0039 Extended Data Figure 4). Collectively, these results suggest that tau_NFT_-induced β-gal gene upregulation did not alter activity at pH 6.0, nor contribute to brain atrophy. Senescent cells exert toxicity to adjacent cells and tissue through the release of toxic secreted molecules. To determine if the NFT-induced SASP-associated profile (Figure 1) affected the surrounding neuronal microenvironment, we performed distance analyses between histologically normal neurons (absence of NFTs), between normal neurons and NFTs, and between NFTs. The distance between neurons was significantly increased in the presence of NFTs (*P* < 0.0001, Figure 3c, f). Specifically, the distance between a non-NFT and NFT, or between two NFTs was greater than two non-NFT neurons (*P* < 0.0001; 1.6-and 1.5-fold, respectively). Collectively, these results led us to conclude that NFTs induced DNA damage, nuclear hypertrophy, and altered the surrounding neuronal microenvironment.

Permanent cell cycle arrest is a hallmark of cellular senescence, and it is induced by DNA damage, oxidative stress and SASP^4^. A recent study reported that DNA damaging agents associated with Parkinson’s disease could induce a senescence-like phenotype in astrocytes^21^. Notably, decreased nuclear lamin B1 expression, the marker used to identify astrocyte senescence *in vivo*^21^, also occurs in AD^22^. In AD and tau transgenic animals, decreased lamin B1 is specific to tau-containing neurons where it causes relaxation of heterochromatin and activates the cell cycle^22^, all of which are features of cellular senescence. Aberrant neuronal cell cycle activity induces AD pathology and neurodegeneration^23^. The cell cycle protein p21, encoded by *Cdkn1a,* is upregulated in many senescent cell types and has been associated with DNA damage during neuronal aging^24^. We found that tau_NFT_ brains expressed 3-fold higher *Cdkn1a* than tau_WT_ mice (*P =* 0.0178, Figure 3g); this was replicated in a separate mouse cohort (*P =* 0.0086, Extended Data Figure 5a). Elevated expression of the cyclin dependent kinase inhibitor 2a, *Cdkn2a,* is one of the most robust markers of cellular senescence, and its protein product, p16^INK4A^, co-localizes with NFTs in human AD^25^. Anti-p16^INK4A^ antibodies are notoriously poor in mouse tissue, so we exclusively measured *Cdk2na* gene expression. Tau_NFT_ *Cdkn2a* forebrain expression was 2.7-and 2.6-fold higher than CTL and tau_WT_, respectively (*P* = 0.0303 and *P* = 0.0352, respectively; Figure 3h); this effect was replicated in an independent mouse cohort (*P* = 0.0016, Extended Data Figure 5b). Similar to mitochondrial dysfunction, genotype-specific changes in *Cdkn2a* were not found in the cerebellum, a brain region without transgenic tau expression or NFTs, indicating that the pathogenic tau microenvironment was required for *Cdkn2a* upregulation (Extended Data Figure 5c). Further, when plotted against brain weight, *Cdkn2a* expression was a strong predictor of brain atrophy across mouse lines (*P* < 0.0001, R^2^ = 0.5615; Figure 3i; and tau_NFT_ mice plotted alone: *P* = 0.0158, R^2^ = 0.5885; Extended Data Figure 5d). These data connect NFTs, *Cdkn2a* and neurodegeneration.

To determine whether senescence-associated *Cdkn2a* expression was linked to NFT density, NFT onset, or merely protein accumulation, we pursued multiple genetic approaches and a pharmacological intervention. Genetic deletion of endogenous mouse tau (microtubule associated protein tau, *Mapt,* knockout) reduced soluble and insoluble tau expression, decreased NFT pathology and neurodegeneration in tau_NFT_ mice^26^ (tau_NFT_-*Mapt^0/0^*). The reduced tau pathology corresponded with 60% lower *Cdkn2a* expression (*P* = 0.0041, Figure 3j), decreased SASP (Extended Data Figure 6) and correlated with decreased brain atrophy (tau_NFT_-*Mapt^0/0^*: 0.4058 ± 0.009 versus age-matched tau_NFT_ *Mapt^wt/wt^*: 0.3451 ± 0.0116; 17.5% difference, *P* = 0.0143). Because tau_NFT_ mice develop aggressive tauopathy with NFT formation in early life, we focused on tau_WT_ mice between 16-28-months-old to detect subtle cellular changes associated with different stages of NFT development and progression. We found that *Cdkn2a* gene expression increased significantly during this age interval, and at 28 months of age tau_WT_ *Cdkn2a* expression was similar to that of 16-month-old tau_NFT_ mice (Extended Data 5e). Concomitantly, at this age, tau_WT_ mice developed NFTs as visualized by Bielschowsky silver staining and immunofluorescence analyses (Extended Data 5f, g). These results provide additional evidence for the association between NFT formation and senescence-associated *Cdkn2a* upregulation. Next to determine if *Cdkn2a* expression was driven specifically by NFTs, or whether AD-associated Aβ protein deposition also increased *Cdkn2a*, we utilized 3xTgAD mice that acquire both AD-associated pathologies with Aβ deposition and NFT onset at 6 and 18-months of age, respectively^27^. In 15-month-old mice with heavy Aβ deposition and phosphorylated tau, but lacking NFT pathology^28^, *Cdkn2a* expression was not elevated (Extended Data Figure 5h) indicating that *Cdkn2a* expression was neither a response to general protein accumulation, nor to pre-NFT tau pathology, but instead required the presence of NFTs. To then determine whether these findings translated to human brains with pure tauopathy (i.e., NFT pathology without other protein aggregates such as Aβ, we acquired human brain tissue with histopathologically confirmed progressive supranuclear palsy (PSP) (Extended Data Table 1 for patient characteristics). PSP is an age-associated tauopathy that clinically manifests as parkinsonism with additional motor abnormalities and cognitive dysfunction^1^, and is neuropathologically defined by accumulation of four-repeat (4R) tau, NFTs, gliosis and neurodegeneration^29^. Consistent with our results from transgenic mice, *CDKN2A* expression correlated with NFT deposition, specifically in the parietal lobe (ANOVA, *P* = 0.0008; Kendall’s Tau rank correlation *P =* 0.059, Figure 3l). Moreover, one individual with the worst cognitive performance, Mini–Mental State Examination (MMSE) score of 12, displayed the highest level of *CDKN2A* expression, and high molecular weight tau (Figure 3m). Collectively, these findings led us to conclude that NFTs were directly linked to senescence-associated *Cdkn2a* upregulation, which in turn was a strong predictor of neurodegeneration and cognitive decline.

Pharmacological agents targeting senescent cells, termed senolytics, have successfully decreased a variety of age-associated pathologies in model organisms, thus are an appealing class of medications to translate to human disease (recently reviewed,^30^). We used the best-characterized senolytics to date, dasatinib and quercetin (DQ), to determine the utility of targeting cellular senescence to treat tau-associated neurodegeneration. To test the effects in late-life we used tau_NFT_ mice on the less aggressive *Mapt^0/0^* genetic background. Beginning at 20-months-old, tau_NFT_-*Mapt^0/0^* and non-transgenic-*Mapt^0/0^* mice were randomized to receive vehicle or DQ at bi-weekly intervals for 3 months. When mice were 23-months-old, brain structure and function were analyzed with MRI and postmortem histopathology (Figure 4). Histological analyses revealed the presence of NFT-containing neurons in both treatment groups. However, DQ treatment reduced senescence burden as evidenced by a 5% decrease in cortical NFTs (*P* < 0.0001; Figure 4a b) and an overall decrease in senescence associated gene expression (*P* = 0.0006; Extended Data Figure 7a-c). Neurons with NFTs in both treatment groups displayed karyomegaly (Figure 4c). This finding is consistent with NFT-containing neurons from tau_NFT_ mice (Figure 3c, d) indicating that NFTs induce this senescence-associated phenotype in multiple transgenic mouse models. Brains from DQ-treated mice displayed a significantly different brain cell specific protein expression profile that was driven by differences in neuronal proteins (NeuN: 25% elevated, *P* = 0.0432 and synaptophysin: 41% elevated, *P* = 0.0416) but not glial or myelin proteins, GFAP or Plp1, respectively (Figure 4d-f). DQ did not alter total human tau protein levels indicating these effects were driven by tau in the form of NFTs (Extended Data Figure 7d, e).

**Figure 4.**
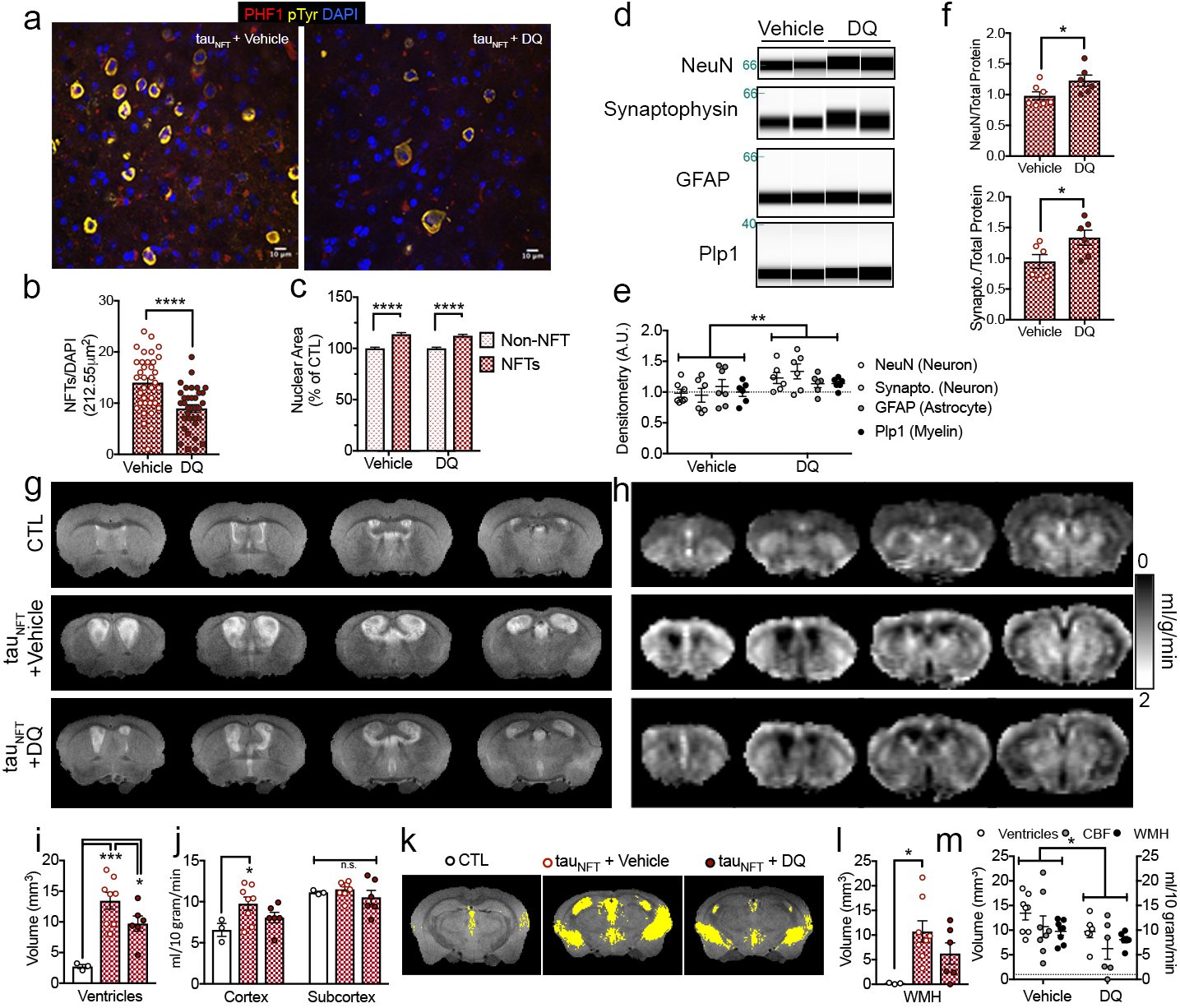
Senolytic treatment reduced NFT burden, ventricular enlargement and cerebral blood flow deficits in 23-month-old tau transgenic mice. (a) Representative brain images analyzed for neurofibrillary tangles in tau transgenic mice treated with either vehicle or dasatinib and quercetin (DQ). (Phosphorylated tau, PHF1, red; total tyrosine phosphorylation, pTyr, yellow; and DAPI nuclei; blue. Scale bar = 10μm). (b) Neurofibrillary tangle counts from n = 3 mice/group sampled from 12 cortical images/mouse and analyzed with unpaired two-tailed t-test, **** *P* < 0.0001. (c) Tau_NFT_ *Mapt^0/0^* neurons with NFTs display karyomegaly; treatment with DQ does not alter DAPI nuclear area of remaining NFTs. Unpaired two-tailed t-test Non-NFT vs. NFT: *P* < 0.0001; Vehicle NFT (n = 369) vs. DQ NFT (n = 429): *P* = 0.4711. Vehicle non-NFT n = 330; DQ non-NFT n = 336. (d) Immunoblot generated by capillary electrophoresis on forebrain homogenates with antibodies against neuron (NeuN and synaptophysin, Synapto.), astrocyte (GFAP) and myelin (Plp1) proteins and (e, f) normalized to total protein. (e) Data analyzed by two-way ANOVA; treatment main effect, *P* = 0.0023; and (f) Unpaired two-tailed t-test, NeuN: * *P =* 0.0432 and Synapto: * *P =* 0.0416. (g) Representative brain images from anatomical T2-weighted MRI and (h) cerebral blood flow MRI. (i) Quantification of ventricle volume analyzed by one-way ANOVA, *P* = 0.0010; Tukey’s *posthoc*: *** *P* = 0.0007, * *P* = 0.0239; Vehicle vs. DQ: Holm-Sidak’s *posthoc* * *P* = 0.05. (j) Cerebral blood flow MRI quantification of cortex and subcortical brain regions among groups (two-way ANOVA brain region main effect, *P* < 0.0001; Tukey’s *posthoc:* * *P* = 0.0253). (k) Representative brain images illustrating areas of white matter hyperintensity (WMH, yellow pixels) and (l) WMH volume analyzed by one-way ANOVA, *P* = 0.0330; Tukey’s *posthoc* * *P* = 0.0291. (m) Composite analysis of tau_NFT_ + vehicle and tau_NFT_ + DQ MRI data analyzed by two-way ANOVA DQ treatment main effect: * *P* = 0.0138. (Tau_NFT_ + vehicle, n=8; tau_NFT_ + DQ, n=6; non-transgenic, n=3; all mice were on a *Mapt^0/0^* background).

The intermittent senolytic dosing schedule allows for rapid drug clearance, instead of chronic activation of molecular targets/pathways common among many therapeutic strategies. To determine if cyclic senescent cell removal produced long-lasting global effects on brain, we used MRI to analyze anatomical structure and cerebral blood flow, a functional defect that occurs in AD and tau_NFT_ mice and is closely associated with cognitive impairment^31^. Pathogenic ventricular enlargement is hallmark of AD, and the rate of ventricular volume change is highly correlated with NFTs^32^. Tau_NFT_ mice recapitulate this pathology on a wild type (Extended Data Figure 1) and *Mapt^0/0^* background (*P* = 0.0007; Figure 4g, i). Though DQ did not completely rescue this pathology to control volume size (tau_NFT_*-Mapt^0/0^* + DQ vs. CTL; *P* = 0.0239, Figure 4i), a finding that is not completely unexpected considering the severity of disease and age of the animals, the treatment tended to reduce ventricular volume pathology compared to vehicle treated tau_NFT_*-Mapt^0/0^* mice (28% decrease, *P =* 0.05, Figure 4i). In cortical brain tissue with tau pathology, cerebral blood flow was elevated in tau_NFT_ *Mapt^0/0^* vehicle-treated mice (48.9%, *P* = 0.0141, Figure 4h, j), which was consistent with previous reports of tau_NFT_ mice on a *Mapt^+/+^* background^31^. DQ abrogated these effects whereby tau_NFT_ mice after treatment were no longer significantly different than non-trasngenic controls (*P* = 0.1527; Figure 4h, j). We also observed white matter hyperintensitites (WMH), areas of increased brightness on T2-weighted MRIs (Figure 4k). The underlying WMH pathology, cerebral small vessel disease, causes chronic ischemia and increased risk of cognitive decline and dementia (reviewed^33^). Unlike the tau_NFT_-*Mapt^0/0^* vehicle treated mice with WMH pathology (*P* = 0.0341; Figure 4k, l), tau_NFT_-*Mapt^0/0^* mice treated with DQ did not display WMH volumes statistically different than control mice (*P* = 0.2458; Figure 4l). Brain atrophy progresses with age in tau_NFT_-*Mapt^0/0^* mice, and was evidenced by a 9.8% decrease in brain volume in tau_NFT_-*Mapt^0/0^* vehicle treated mice compared to CTLs (*P* = 0.0026, Extended Data Figure 8). However, a significant difference in brain volume between CTLs and tau_NFT_ *Mapt^0/0^* mice that received DQ was not observed. Closer evaluation of brain regions revealed that this effect was driven by cortical atrophy. While subcortical brain region volume did not differ among groups, cortical volume of vehicle treated tau_NFT_-*Mapt^0/0^* mice was 31% lower than controls; DQ mitigated this effect (*P* = 0.0092 and *P* = 0.0274, vehicle and DQ, respectively; Extended Data 8a, b). Overall, the composite effect of DQ treatment in tau_NFT_ mice was a significant global benefit (*P* = 0.0138; Figure 4m); targeting cellular senescence reduced total NFT burden and SASP expression, enhanced neuronal protein expression, and benefited overall tissue structure and cerebral blood flow in a late-life, advanced stage AD mouse model.

The inability to effectively treat tau-associated diseases arises, in part, from a limited understanding of processes driving neurodegeneration during the prodromal period. We have identified cellular senescence, the quintessence of latent tissue degeneration, as a cellular mechanism upregulated in tau-associated neurodegeneration. Findings in NFT-developing transgenic mice, postmortem human AD and PSP brain tissue support this concept. Pathogenic tau induces a traditional neuroinflammatory response by activating microglia and astrocytes (for recent review^34^). Our data suggest that NFT-containing neurons may be active participants in perpetuating the inflammatory response as well. Through gene expression of microdissected neurons, assessment of neuronal nuclei and toxicity to adjacent tissue, and clearance with senolytics, our data suggest that NFT-containing neurons acquire a senescence-like phenotype. In this way, NFTs formed in early pathogenic stages may contribute to neurodegeneration later in life through senescence-like mechanisms by altering the bioenergetic state of the brain and upregulating the toxic SASP in various cell types, including neurons. Moreover, therapeutically targeting cellular senescence effectively interrupted this chronic neurodegenerative cascade to improve tau pathology, brain structure and cerebral blood flow deficits even in the presence of established tau pathology in a late-life advanced disease state.

## METHODS

### Mice

All animal experiments were carried out following National Institutes of Health and University of Texas Health Science Center at San Antonio (UTHSCSA) Institutional Animal Care and Use Committee guidelines. We used 16 to 32-month-old male and female rTg4510 and rTg21221 mice that reversibly express P301L mutant human tau or wild type human tau 4R02, respectively, on either a wild type or *Mapt* knockout Bl6/FVB genetic background^9,10,26^. Non-transgene expressing littermates from rTg4510 and rTg21221 are used as controls; since no differences were found between these control lines, only littermates from rTg4510 are used here (Extended Data Figure 1f-i). The mice were bred by Rose Pitstick and George A. Carlson at McLaughlin Research Institute, Great Falls, MT. Mouse euthanasia, brain dissection and preparation was performed as previously described^18,28^.

### Ingenuity Pathway Analyses (IPA)

The GEO accessions GDS2795 and GSE56772 were accessed from the GEO Profiles database^3,8^ with Rstudio version 1.0.143. Because the GDS2795 datset was a within subjects design, ratios of NFT vs CTL gene expression were generated for each subject. Mean ratios of each gene from GDS2795 and fold change values from GSE56772 were uploaded into IPA software (IPA, QIAGEN Inc.,https://www.qiagenbioinformatics.com/products/ingenuity-pathway-analysis). For GDS2795 GeneBank Accession IDs were used and 34910 out of 54675 genes were identified by IPA software. The expression fold change cutoff value was set at 3 (both down/upregulated) compressing the analyses to 3048 genes. For GSE56772 LIMMA package was used to determine fold change and p-values. The *P*-value cutoff for IPA analysis was set at *P* < 0.01 yielding 1294 transcripts, 738 down-and 556 up-regulated. We utilized IPA causal analytic tools^35^ to elucidate predicted upstream regulators, as well as disease and biological functions with significant z-scores enriched in our data set.

Findings from GDS2795 were replicated with more stringent criteria (p < 0.05 for NFT/CTL ratios with no fold change limit) allowing for 1715 genes to be uploaded into IPA with similar results. Similarly, these findings were replicated a third time using the LIMMA package, the most common method for microarray analysis. This method did not take into account within subjects design. Using a *P* < 0.05, 1219 differentially regulated genes were uploaded for IPA analyses; the results were similar to the original findings. Furthermore, results from GSE56772 were replicated using Gene Set Enrichment Analysis (GSEA) with default setting and similar results were obtained.

### RNA extraction and qPCR

Frozen forebrain and cerebellum were powdered in liquid nitrogen. RNA was extracted from ~25mg of each respective brain (or brain region) using the RNAqueous 4PCR^®^ kit (Ambion), following the manufacturer protocol including the 15 minute DNase treatment. qPCR was performed on 25ng RNA using the TaqMan^®^ RNA-to-CT™ 1-step kit. All gene expression analyses were made using Taqman gene expression assays. RNA Polymerase II Subunit J (Polr2j) expression was used as an internal control for both mouse and human gene expression assays, Mm00448649_m1 and Hs01558819_m1, respectively. Taqman genes expression identifiers for target genes are as follows: mouse and human *Cdkn2a*: Mm00494449_m1 and Hs00923894_m1, respectively; other mouse genes: *Cdkn1a:* Mm00432448_m1*; Glb1:* Mm01259108_m1*; Cxcl1:* Mm04207460_m1; *Tlr4*: Rn00569848_m1; *Il1β*: Mm00434228_m1; *TNF*: Mm00443258_m1. The senescent cell population comprises a small proportion of all cells in a tissue; therefore SASP gene expression values were normalized to neuronal *Mapt,* the senescence-susceptible neuronal population. qPCR was performed using the Applied Biosystems 7900HT Sequence Detection System, with SDS software version 2.3. Cycle profile was performed using the kit manufacturer protocol.

### Protein extraction and capillary electrophoresis

50mg frozen forebrain was used for subcellular fractionation and capillary electrophoresis as previously described^28,36^. Briefly, frozen tissue was powdered in liquid nitrogen, then homogenized with dounce and pestle and fractionated following manufacture protocol (Subcellular Protein Fractionation Kit, ThermoFisher Scientific, USA). Protein concentrations were determined with BCA (Biorad, USA); 2ug protein was used for capillary electrophoresis. Antibodies were diluted in Wes Antibody Diluent to the final working concentrations: p65, 1:50 (Cell Signaling, D14E12; Beverly, MA, USA); phospho-Ser139 H2A.X, 1:50 (Cell Signaling, 20E3; Beverly, MA, USA); HT7, 1:1000 (Pierce/Invitrogen, USA); NeuN, 1:50 (Millipore, MAB377; Temecula, CA, USA): GFAP 1:200 (Cell Signaling, D1F4Q; Beverly, MA, USA); synaptophysin, 1:50 (Cell Signaling, D35E4; Beverly, MA, USA); Plp1, 1:100 (Sigma, HPA004128; St. Louis, MO, USA). See Extended Data Table 2 for complete antibody information. Protein quantification is performed by normalizing to total protein concentration^37,38^; (Extended Data Figure 9).

### Histology

Brains were fixed in 4% PFA for 48 hours, transferred to PBS containing 0.02% sodium azide and vibratome sectioned at 30μm. Sections were washed 3x with TBS (pH 7.4), and incubated in 50% ethanol for 5 mins; followed by 70% ethanol for 5 minutes. The sections were then submerged in 0.7% sudan black b dissolved in 70% ethanol for 5 minutes to quench lipofuscin-like autofluorescence. Tissues were then rinsed 3 times for 1-2 min in 50% ethanol. Following this step, tissue sections were transferred from 50% ethanol to TBS and proceeded to immunofluorescence staining as described previously^28,36^. Primary antibodies used: PHF1 (1:100, kind gift from Dr. Peter Davies), NeuN (1: 500 Cell Signaling, D3S31; Beverly, MA, USA), Histone 3 (1:400, Cell Signaling, D1H2; Beverly, MA, USA) (Extended Data Table 2). Secondary antibodies: Goat anti-Mouse IgG (H+L), Alexa Fluor 594 and Goat anti-Rabbit IgG (H+L), Alexa Fluor 488 (1:1500, Thermo Fisher Scientific). Imaging was performed using a Zeiss LSM 780 confocal microscope, with ZEN 2.3 software.

#### Confocal Image Analyses

Image analyses were conducted using ImageJ’s FIJI. Analyses were performed on confocal z-stacks imaged at 40x magnification. A maximum intensity image was created by compressing four z-stack planes. For nuclei size analyses, using DAPI, cell nuclei were identified and thresholded for proper nucleus selection. All analyzed DAPI fields were applied a bandpass filter under the same conditions, applied a threshold, and measured using particle analysis excluding particles smaller than 25 μm^2^. All particles measured in the analysis were checked for mislabels and any particles that included 2 nuclei or exhibited abnormal/incorrect selection were excluded from analysis. Cell type was identified using NeuN (neurons) and PHF1 (NFT-bearing neurons) immunofluorescence. For within-subject analysis of NFT-bearing compared to healthy neurons, cell area data were taken and averaged from 16 separate fields (Z-projections) across rTg4510 mice (n=3). NFT-bearing neurons were paired with a nearest-neighbor neuron (determined by NeuN positivity) for paired statistical analysis. For distance analyses, all neurons (including NFTs) were outlined and number identified by thresholding on the NeuN channel. Using the line tool, the distance from NFT-bearing neurons to each of their nearest neighbors (NFTs and non-NFTs) was measured and recorded. As a control, the distance from randomly selected healthy neurons to each of their neighboring neurons (non-NFTs) was also measured.

#### SA β-gal Staining

Following euthanasia, brains were immediately removed and fresh-frozen in an isopentane/liquid nitrogen slurry. The frozen brains were immediately adhered to the cryotome chuck with optimal cutting temperature compound (OCT) pre-cooled to −18°C; 10μm coronal sections were collected, and mounted on superfrost plus microscope slides (FisherScientific). After sectioning, slides were fixed for 10 minutes in 2% paraformaldehyde/0.2% glutaraldehyde at room temperature, rinsed 3x in TBS and stained with SA β-gal staining solution overnight^17^. Following SA β-gal staining, sections were processed for immunofluorescence as described above.

#### Brightfield/SA β-gal Counts

SA β-gal brightfield images were taken on a Nikon Eclipse Ci-L microscope, with a digital site DS-U2 camera (NIS-Elements software BR 4.51.00). Coronally sectioned mouse brains (10μm) were evaluated using DAPI, SA β-gal positive and negative cells in the CA2 region of the hippocampus were counted across 8 tissue sections per animal (n=5 per genotype). Staining was considered positive when granules of blue stain were present.

### High Resolution Respirometry

HRR was conducted using two Oxygraph-2k (model D & G) machines from Oroboros Instruments (Austria). To minimize mitochondrial damage associated with mitochondrial isolation techniques, we measured oxygen consumption in fresh brain tissue homogenates^39^. Whole hippocampus, cortex and cerebellum were homogenized with ~15 strokes using a Kontes glass homogenizer in 5% w/v ice cold MiRO6. Two mg of brain homogenate were loaded into the chamber and experiments were carried out when oxygen concentration in each well was saturated under atmospheric conditions (~190nM/mL O_2_). All reagents and SUIT protocol were described previously^40^ with small modifications. Briefly, 1.25mM ADP was sufficient for saturation in brain homogenate, rotenone was added at a concentration of 1μM, and FCCP was added in a single injection at a concentration of 0.5μM.

#### DQ

Control and tau_NFT_ *Mapt*^0/0^ mice aged 19-20 months were randomized to receive DQ senolytic (5mg/kg dasatinib (LC Laboratories, Woburn, MA) with 50mg/kg quercetin (Sigma-Aldrich, St. Louis, MO)) or vehicle (60% Phosal 50 PG, 30% PEG 400, and 10% ethanol) via oral gavage as described previously^41^. Mice were weighed and fasted for 2 hours prior to treatment. One month after the first treatment, senolytic or vehicle gavage continued on a bi-weekly basis for a total of six treatment sessions over twelve weeks. Within two weeks of the final treatment, all mice underwent MRI analyses.

#### MRI

MRI experiments were performed on an 11.7 Tesla scanner (Biospec, Bruker, Billerica, MA). A surface coil was used for brain imaging and a heart coil^42^ for arterial-spin labeling. Coil-to-coil electromagnetic interaction was actively decoupled. Mice were maintained on 1.5% isoflurane anesthesia for MRI duration. *Anatomical MRI*: Anatomical images were obtained using a fast spin echo sequence with a matrix = 128×128, field of view (FOV) = 1.28cmx1.28 cm, repetition time (TR) = 4000 ms, effective echo times = 25 ms. Thirty 1-mm coronal images were acquired with 4 averages. Total scan time= 8.5 minutes. *Cerebral blood flow (CBF)* was measured using continuous arterial spin labeling technique with single shot, spin-echo, echo-planar imaging and analyzed as previously described^43^. The images were acquired with partial Fourier (3/4) acquisition, matrix size = 64×64, FOV = 1.28cmx1.28 cm, TR = 3000 ms, TE = 10.59 ms, post labeling delay=350ms. Seven 1-mm coronal images were acquired with 100 repetitions. Total scan time=10 mins. *Analyses:* MRI analysis was conducted using Stimulate (Center for Magnetic Resonance Research, University of Minnesota Medical School, Minneapolis, MN) running on a CentOS5 Linux Operating System^44^. Anatomical MRI images were used to measure cortex, subcortex, ventricle, white matter hyperintensities (WMH) and whole brain volume. The desired Region of Interest (ROIs) were outlined and volumes were obtained by multiplying ROI total voxels by voxel volume (0.004 mm^3^). Ventricle and WMH volume was obtained by thresholding anatomical image voxels to highlight regions of greater intensity, followed by ROI traces of the target regions. Whole brain volume was obtained by ROI trace after removal of the skull using a local Gaussian distribution 3D segmentation MATLAB code^45^.

### Statistical Measures

#### Transgenic mouse analyses

Measurements were taken from distinct samples. Key finding were repeated in separate mouse cohorts and are listed in Extended Data Figures. Each age cohort for all analyses contained 3–9 animals (specified in figure legends); both males and females were included. Only females were included in DQ treatment and MRI analyses due to animal availability. Statistics were not used to predetermine sample sizes, but instead was determined empirically from previous experimental experience with similar assays, and/or from sizes generally employed in the field. Data are expressed as mean ± standard error of the mean (s.e.m.). Genotype comparisons were analyzed with one-way analysis of variance (ANOVA) with Tukey post-hoc analysis unless stated otherwise. Respirometric data, brain volume and cerebral blood flow data were analyzed using two-way ANOVAs (genotype x respirometric parameter) and (treatment x brain region), respectively, with Tukey’s post hoc comparisons.

#### Human PSP brain tissue analyses

the PSP group contained 14 samples and were compared to 10 age-matched controls with both sexes included; significance was determined with unpaired two-tailed t-test. We performed ANOVA analysis of the log base 10 transformed CDKN2A expression as predicted by the levels of AD pathology, in addition we used the more conservative Kendall’s Tau rank correlation to test this association as well. The human CDKN2A and AD pathology analyses were performed using the R v3+ (Vienna, Austria) environment for statistical computing using an accountable data analysis process. All other data were analyzed using GraphPad Prism version 7.0c for Mac OS X, GraphPad Software, San Diego California, USA, www.graphpad.com/. Data were considered statistically different at *P* < 0.05.

## Acknowledgements

We thank Drs. You Zhou for independently performing IPA analyses to confirm the results presented here; Jia Nie for assistance with oral gavage; Ning Zang for assistance collecting tissues; and Ji Li for maintaining the mouse colony, assistance with tissue collection and laboratory management. We thank Mr. Anthony Andrade for providing technical assistance with sectioning mouse brains. The Nathan Shock Pathology Core and Nathan Shock Metabolism Core provided the cryostat and Oxygraph-2k, respectively. Drs. Yuji Ikeno and Judith Campisi provided technical advice on SA β-gal staining. We would like to acknowledge Dr. Jonathan Gelfond for assisting with the data analysis, and was supported by NIH grants NIA Shock Center P30AG013319 and NIA Pepper Center P30AG044271. The authors acknowledge Karen H. Ashe for development of the rTg4510 mouse line. We thank the McLaughlin Research Institute: Dr. George Carlson and Rose Pitstick for breeding all mouse lines used here, and Dr. Andrea Grindeland for performing silver staining. We are grateful to the Banner Sun Health Research Institute Brain and Body Donation Program of Sun City, Arizona for the provision of human brain tissue. The Brain and Body Donation Program is supported by the National Institute of Neurological Disorders and Stroke (U24 NS072026 National Brain and Tissue Resource for Parkinson’s Disease and Related Disorders, the National Institute on Aging (P30 AG19610 Arizona Alzheimer’s Disease Core Center), the Arizona Department of Health Services (contract 211002, Arizona Alzheimer’s Research Center), the Arizona Biomedical Research Commission (contracts 4001, 0011, 05-901 and 0110 for the Arizona Parkinson’s Disease Consortium) and the Michael J. Fox Foundation for Parkinson’s Research. This work was supported by the San Antonio Nathan Shock Center for Excellence, the UT Health Science Center School of Medicine Briscoe Women’s Health and the US Department of Veterans Affairs Career Development Award (IK2BX003804) awarded to M.E.O. N.M. is supported by R01-DK80157, R01-DK089229, P30 AG013319 (San Antonio Nathan Shock Center), and P30 AG044271 (San Antonio Claude D. Pepper Older Americans Independence Center). J.M.V. is supported by a Biology of Aging T32 Training Grant (T32 AG021890).

## Author Contributions

N.M. provided guidance and support to authors and edited the manuscript. J.M.V. designed and conducted IPA, GSEA and respirometry experiments, analyzed and interpreted data, prepared figures, wrote methods, independently confirmed key results of gene expression and histological measures, helped write and edit the manuscript. K.S. managed senolytic treatment experiments, performed histological staining, conducted blinded histological analyses of immunofluorescence data, wrote methods and edited the manuscript. E.B. assisted with senolytic treatments, conducted blinded analyses of anatomical MRI images and wrote corresponding methods. Q.S. performed all MRI experiments and analyses, conducted blinded analyses of CBF data and provided oversight to E.B. in anatomical analyses. C.S.T. performed histology, SA β-gal staining and blinded quantification, developed FIJI analysis protocols, wrote methods and edited the manuscript. A.Z. conducted blinded histological analyses of immunofluorescence data, wrote methods and edited the manuscript. M.E.O. conceived, supervised and attained funding for the project; designed the experiments, conducted experiments, analyzed and interpreted data, provided guidance and supervision to co-authors, prepared figures and wrote the manuscript.

## Author Information

Reprints and permissions information is available at www.nature.com/reprints. The authors declare no competing financial interests. Readers are welcome to comment on the online version of the paper. Publisher’s note: Springer Nature remains neutral with regard to jurisdictional claims in published maps and institutional affiliations. Correspondence and requests for materials should be addressed to MEO.

## Online Content

Extended Data display items are available in the online version of the paper; references unique to these sections appear only in the online paper.

**Supplementary Information** is available in the online version of the paper.

## Data Availability Statement

References for source data for Figure 1 are provided with the paper; data that support Figure 1 findings are available from the corresponding author upon reasonable request. All other data supporting the findings of this study are available within the paper, and its extended data files.

---

**Extended Data Figure 1.**
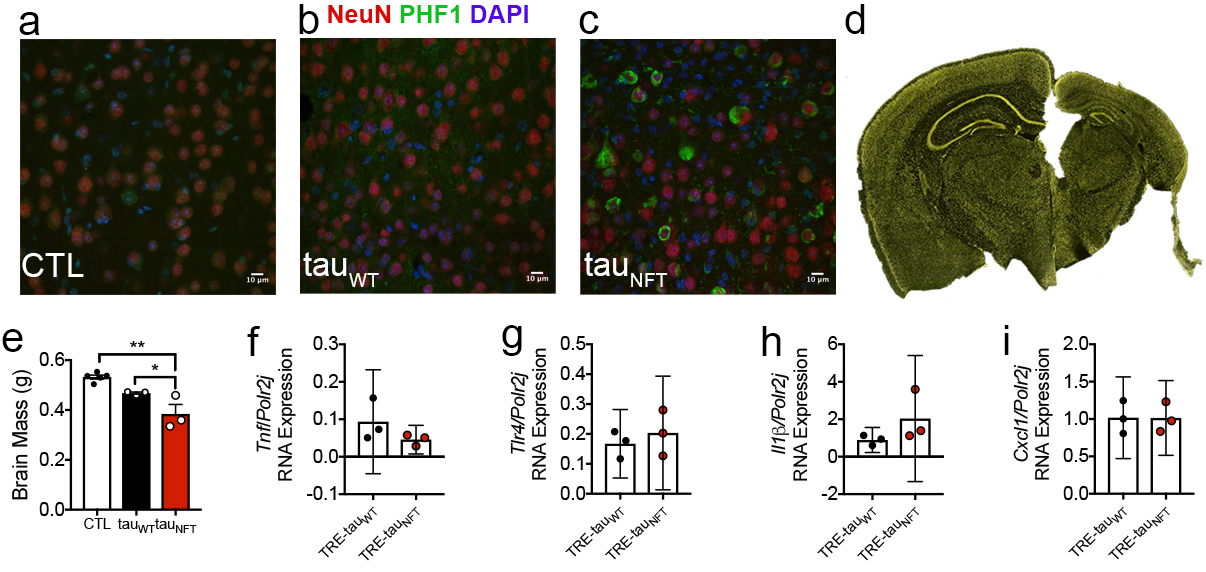
The tau_NFT_ mouse model develops neurofibrillary tangles and brain atrophy. (a-c) Histological staining of cortical brain sections from 16-month-old (a) CTL, (b) tau_WT_ and (c) tau_NFT_ mice with antibodies against neurons (NeuN, red); neurofibrillary tangles (PHF1, green) and nuclei (DAPI, blue). Only tau_NFT_ mice develop neurofibrillary tangles at this age. Scale bar 10μm. (d) Histological comparison of CTL hemibrain (left) to tau_NFT_ mouse hemibrain (right). Images illustrate severe forebrain atrophy with ventricular enlargement in tau_NFT_ mice. (e) Brain weight measurements reflect the significant neurodegeneration in tau_NFT_ mice (n=3/group; one-way ANOVA: *P* = 0.0015, Tukey’s posthoc analysis: CTL vs. tau_WT_ = 0.0644, CTL vs. tau_NFT_ = 0.0011; tau_WT_ vs. tau_NFT_ = 0.0477. Data are graphically represented as mean ± s.e.m. (f-i) Quantitative gene expression analyses on RNA isolated from CTL non-transgene expressing littermates of tau_WT_ mice (TRE-tau_WT_: closed circles) and tau_NFT_ mice (TRE-tau_NFT_: red circles) mouse forebrain indicates statistically similar expression by unpaired two-tailed t-test. (f) *Tnfa*: *P* = 0.2256 (g) *Tlr4*: *P* = 0.5230 (h) *Il1b*: *P* = 0.2244 and (i) *Cxcl1: P* = 0.9808. Data are graphically represented as mean with 95% CI.

**Extended Data Figure 2.**
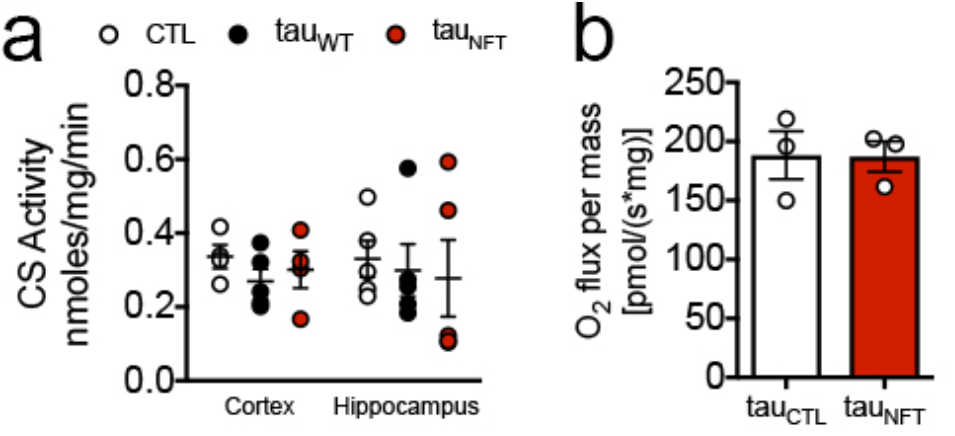
Mitochondrial dysfunction was not due to decreased mitochondrial content, but instead dependent on neurofibrillary tangle pathology. (a) Biochemical analyses indicated that tau_NFT_ mice had comparable citrate synthase (CS) activity in cortex and hippocampus (n=5/group). (b) Moreover, total oxygen consumption in the cerebellum, a brain region devoid of tau pathology did not differ between genotypes. CS activity: one-way ANOVA: (cortex: *P =* 0.7452; hippocampus: *P =* 0.9468; n=3/group). Cerebellum oxygen consumption: Unpaired two-tailed t-test: *P =* 0.9671. Data are graphically represented mean ± s.e.m.

**Extended Data Figure 3.**
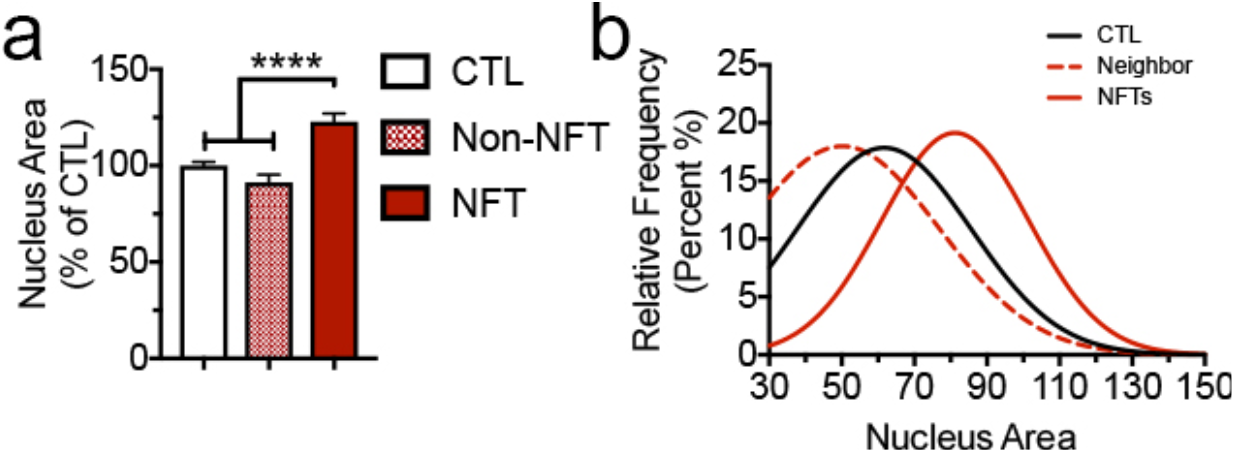
Neurofibrillary tangles induced neuronal karyomegaly. Cortical brain sections were immunostained, and neuronal nuclei were measured. The anti-histone 3 antibody was used to identify nuclear border, anti-HT7 to identify human tau and DAPI to identify cell nuclei. (a). Quantification of histone 3 area. One-way ANOVA: *P* < 0.0001. Tukey’s *post hoc* CTL vs. Non-NFT neighbor, *P* = 0.1359; CTL vs. NFT, *P* < 0.0001; NFT vs non-NFT neighbor, *P* < 0.0001. Brains from n=3 CTL and n=3 tau_NFT_ mice were analyzed. (b) A Gaussian distribution derived from the frequency of cells (y-axis) plotted against the nuclei area in μm^2^ (14 frequency bins generated for each neuronal nuclei category n = 272 NFT neurons counted, n = 272 non-NFT neighbor cells counted; n = 61 control cells). Data are graphically represented as mean ± s.e.m.

**Extended Data Figure 4.**
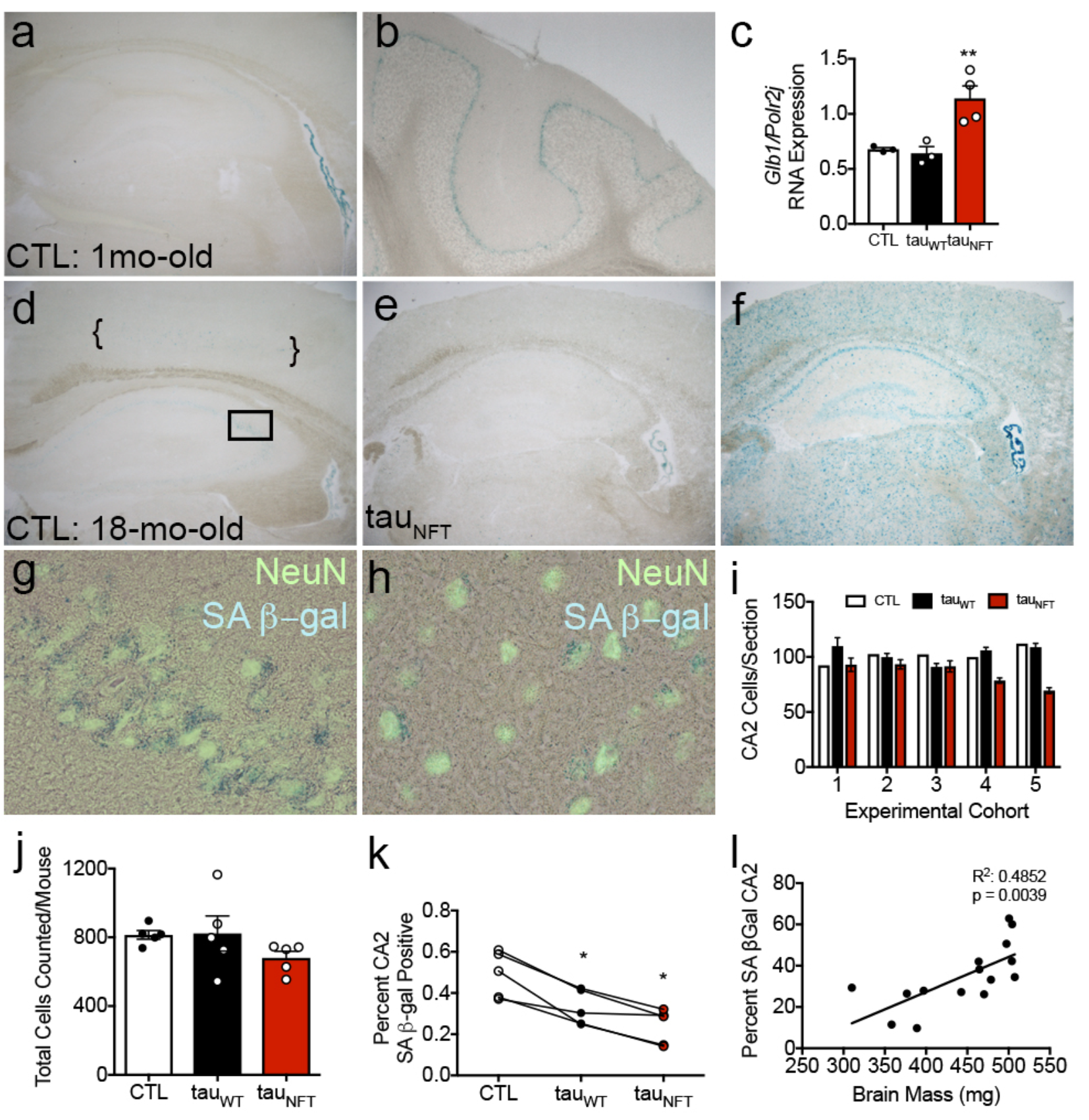
Senescence associated beta galactosidase reactivity was positively correlated with brain mass, but not neurofibrillary tangle pathology. (a) Senescence associated beta galactosidase (SA β-gal) staining carried out at pH 6.0 illustrated reactivity in 1-mo-old control mouse forebrain and (b) cerebellum. (c) Quantitative gene expression analyses on RNA isolated from CTL (open bar), tau_WT_ (closed bar) and tau_NFT_ (red bar) mouse forebrain revealed significantly upregulated lysosomal hydrolase *Glb1* gene expression in tau_NFT_ mice; ANOVA, *P* = 0.0070; CTL: n=3; tau_WT_ n=3; tau_NFT_: n=4. (d) SA β-gal staining of control 18-mo-old brain; CA1 (box) and cortex (brackets). (e) Tau_NFT_ mouse brains stained with β-galactosidase developed very low reactivity at pH 6.0 but (f) positive β-galactosidase activity was observed in tau_NFT_ mice when stained at physiological pH 4. (g-h) Immunostaining with anti-NeuN revealed co-labeling of SA β-gal positive cells with neurons. (i) Five experimental cohorts of CTL, tau_WT_ and tau_NFT_ 18-mo-old mice were analyzed for SA β-gal reactivity at pH 6.0; both males and females were included. The number of CA2 cells sampled on each histological slide was plotted; (j) the total number of CA2 cells counted did not differ among genotypes. (k) The percentage of SA β-gal positive CA2 cells was significantly lower in tau_WT_ mice than controls, and tau_NFT_ mice contained significantly fewer SA β-gal positive CA2 cells than CTL and tau_WT_ mice. (Repeated measures one-way ANOVA: *P* = 0.0049). (l) The percentage of SA β-gal positive CA2 cells was significantly correlated with brain mass (R^2^ = 0.4852, *P* = 0.0 039). Data are graphically represented as mean ± s.e.m.

**Extended Data Figure 4.**
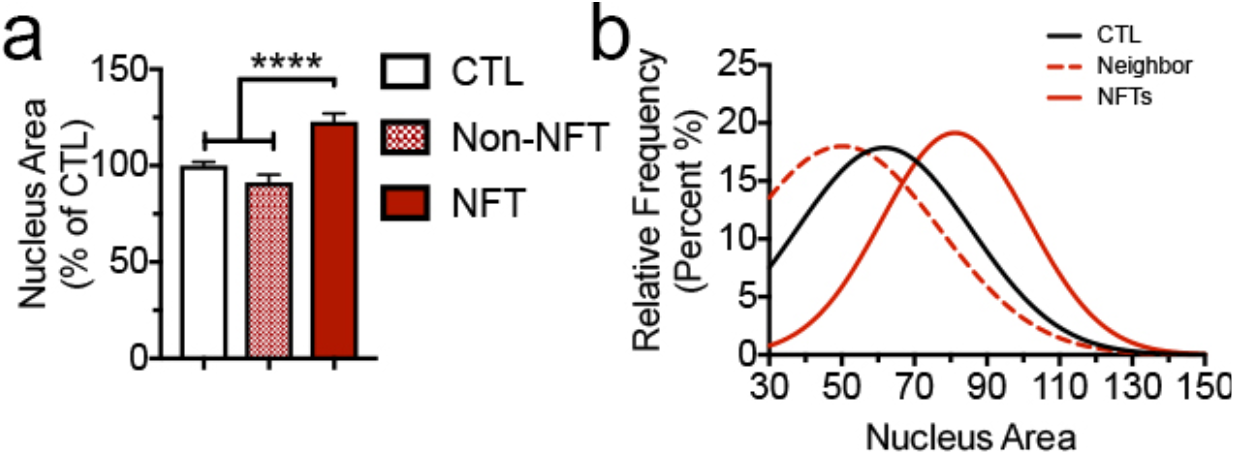
Neurofibrillary tangles induced neuronal karyomegaly. Cortical brain sections were immunostained, and neuronal nuclei were measured. The anti-histone 3 antibody was used to identify nuclear border, anti-HT7 to identify human tau and DAPI to identify cell nuclei. (a). Quantification of histone 3 area. One-way ANOVA: *P* < 0.0001. Tukey’s *post hoc* CTL vs. Non-NFT neighbor, *P* = 0.1359; CTL vs. NFT, *P* < 0.0001; NFT vs non-NFT neighbor, *P* < 0.0001. Brains from n=3 CTL and n=3 tau_NFT_ mice were analyzed. (b) A Gaussian distribution derived from the frequency of cells (y-axis) plotted against the nuclei area in μm^2^ (14 frequency bins generated for each neuronal nuclei category n = 272 NFT neurons counted, n = 272 non-NFT neighbor cells counted; n = 61 control cells). Data are graphically represented as mean ± s.e.m.

**Extended Data Figure 5.**
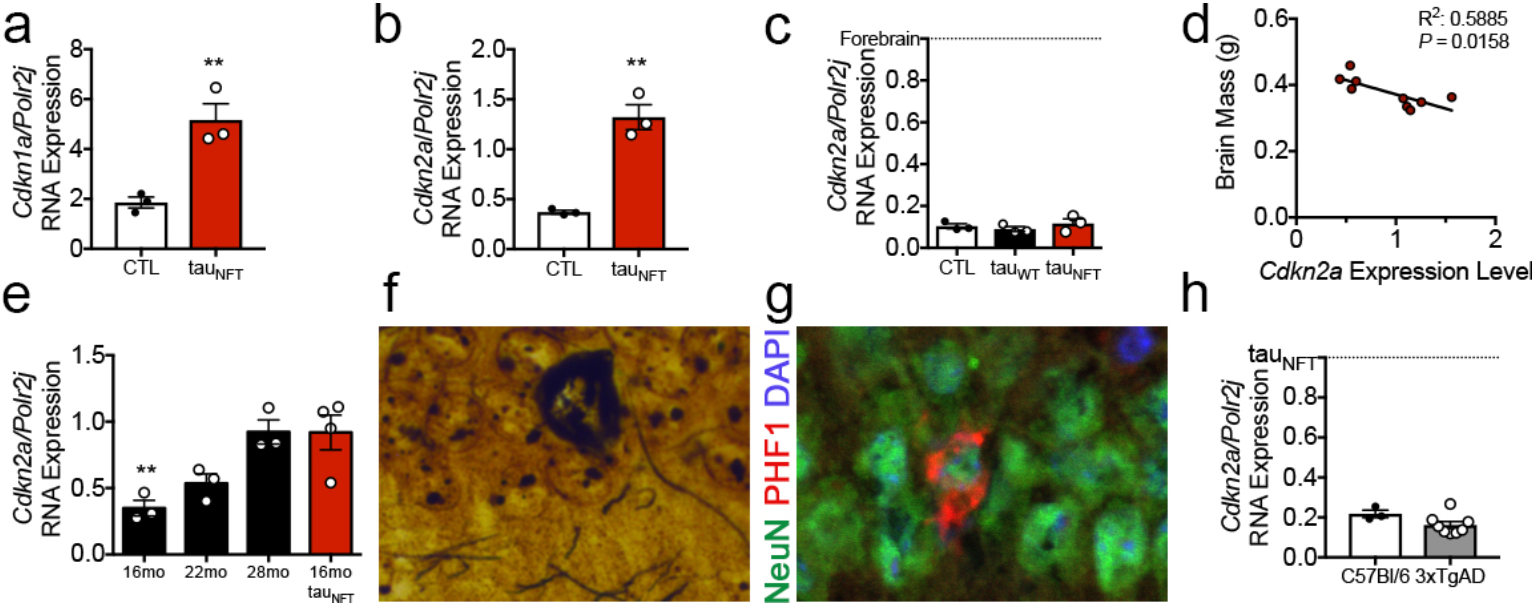
Senescence-associated gene expression was linked to neurofibrillary tangle formation. (a) To test reproducibility of *Cdkn1a* and (b) *Cdkn2a* gene expression findings, we repeated quantitative gene expression analyses on a separate mouse cohort. Unpaired two-tailed t-test: *Cdkn1a*: *P* = 0.0086 and (b) *Cdkn2a*: *P* = 0.0016; tau_NFT_ mice (closed bars) and CTLs (TRE-tau_NFT_: open bars); n=3/group). (c) No differences in *Cdkn2a* expression were found in the cerebellum among genotypes (*P* = 0.7006); all express significantly less *Cdkn2a* than tau_NFT_ mouse forebrain set to y=1, *P* < 0.0001; n=3/group. (d) Total brain mass in tau_NFT_ mice was inversely correlated with *Cdkn2a* expression (n=9; R^2^ = 0.5885, *P =* 0.0158). (e) Tracking *Cdkn2a* expression in tau_WT_ mice revealed a significant age-dependent increase. *P* = 0.0043; n = 3/group for tau_WT_ and n=4 tau_P301L_. In contrast to significantly lower expression than tau_NFT_ mice at 16-mo-old (*P* = 0.0075), Dunnett’s multiple comparison test indicates that at 22-mo-old, tau_WT_ mouse *Cdkn2a* expression was no longer statistically lower than tau_NFT_ mice (*P* = 0.0577) and by 28-mo-old they were statistically the same (*P* = 0.999). (f) Bielschowsky silver staining and (g) immunofluorescence both revealed neurofibrillary tangles in 18-mo-old tau_WT_ mouse hippocampal CA1 (NeuN, neuron, green; PHF1: phosphorylated tau, red; DAPI, blue, nuclei). (h) qPCR analyses of RNA extracted from 3xTgAD mice with Aβ plaques was compared to tau_NFT_ set at y = 1. 3xTgAD *Cdkn2a* expression was no different than age-matched C57BL/6 mice. Unpaired two-tailed t-test, *P* = 0.1081; n = 3 WT, n = 6 3xTgAD. Both mouse cohorts expressed significantly less *Cdkn2a* than tau_NFT_ mice (ANOVA, *P* < 0.0001). Mo = months. Data are graphically represented as mean ± s.e.m.

**Extended Data Figure 6.**
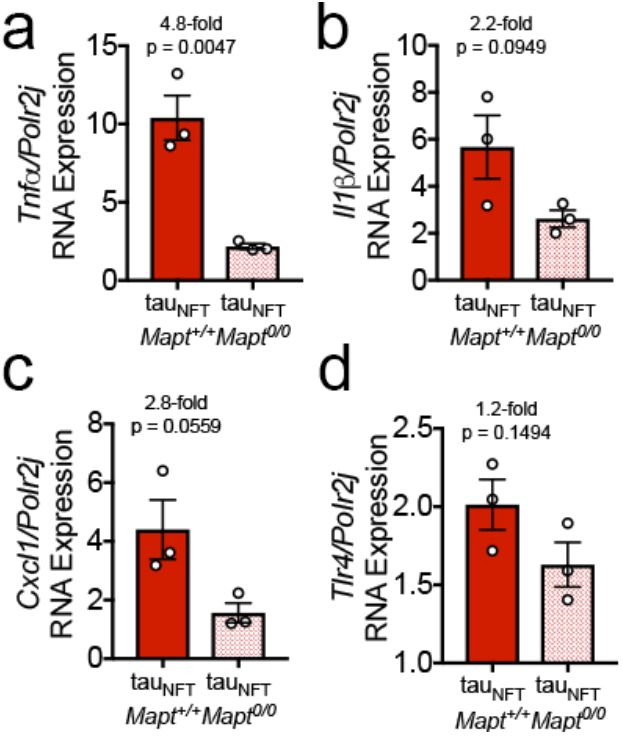
Genetically ablating endogenous mouse tau reduced production of SASP. Quantitative gene expression analyses on RNA isolated from tau_NFT_ mice on a *Mapt* wild type background (closed bars) and *Mapt* knockout background (hatched bars) mouse forebrain revealed a reduction in (a) *Tnfa*, *P =* 0.0047; (b) *Il1b, P =* 0.0949; (c) *Cxcl1, P =* 0.0559; and (d) *Tlr4, P =* 0.1494. N=3/group were analyzed by unpaired two-tailed t-test. Data are graphically represented as mean ± s.e.m.

**Extended Data Figure 7.**
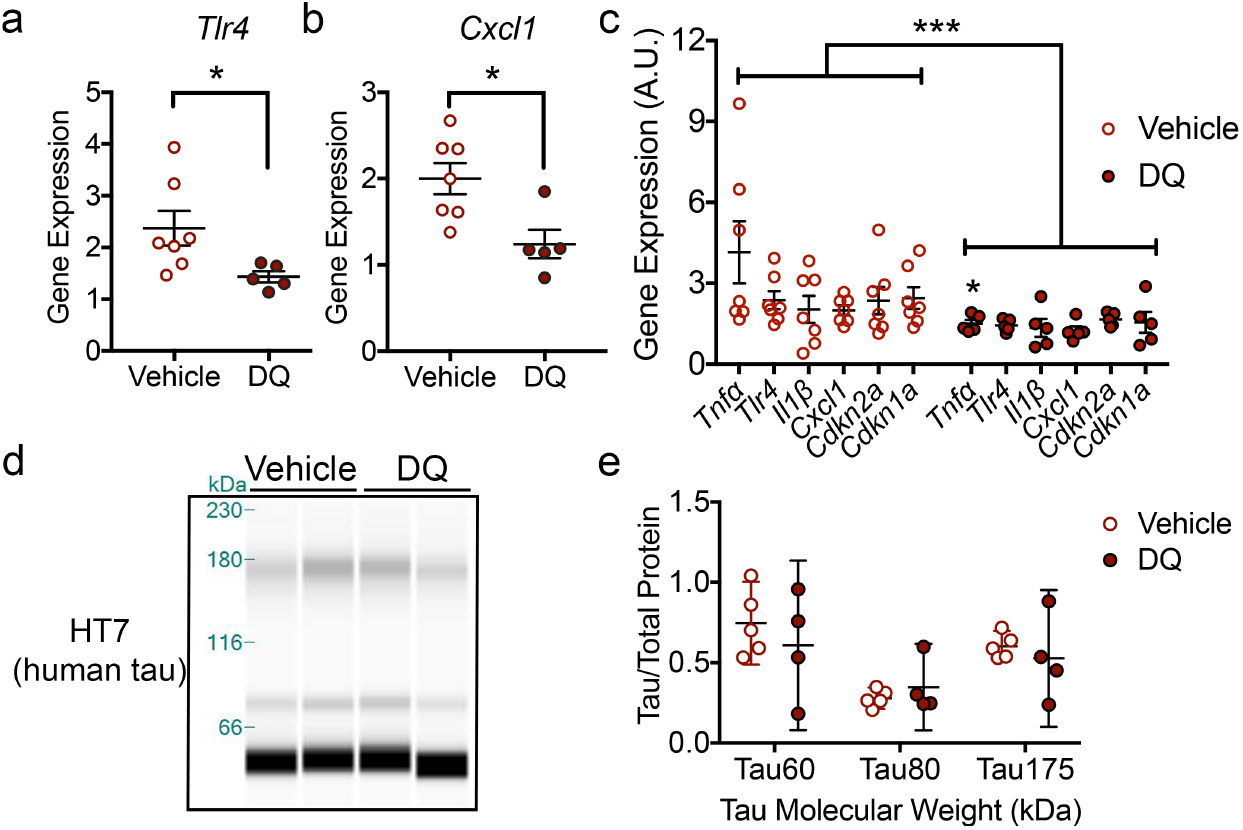
Senolytic treatment decreased senescence-associated gene expression, but did not alter total tau protein levels. (a-c) Quantitative gene expression analyses on RNA isolated from vehicle-treated (open symbols, n=7) and DQ-treated (red closed symbols, n = 5) hippocampal tissue. *Tlr4* and *Cxcl1* gene expression were significantly different by unpaired two-tailed t-test, *: *P* = 0.0459 and *: *P* = 0.0142, respectively. (c) Composite comparison of senescence associated genes between vehicle and DQ treatment analyzed with two-way ANOVA. DQ treatment main effect, \*\*\**P* = 0.0006; Tukey’s *posthoc,* * *P* = 0.0325. Gene expression values were normalized to the *Mapt* gene level. (d) Representative capillary electrophoresis immunoblot using total human tau antibody (HT7) on brain homogenates from vehicle or senolytic (dasatinib and quercetin, DQ) treated mice. (b) Densitometric normalization of total tau immunoreactivity against total protein indicates similar protein expression between treatment groups. Vehicle, n=5; DQ, n=4. Data are graphically represented as mean with 95% CI.

**Extended Data Figure 8.**
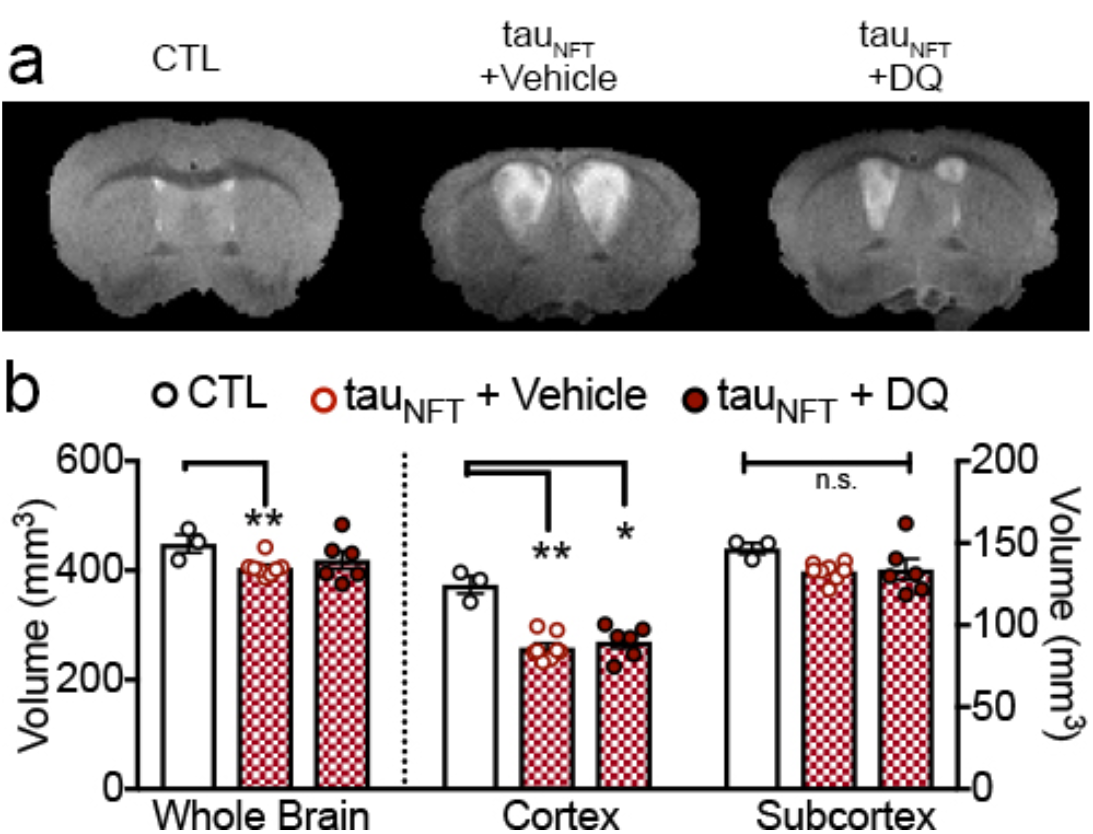
Senolytic treatment had modest effects on total brain volume. (a) Anatomical T2-weighted MRI of tau_NFT_ *Mapt^0/0^* mice (b) Whole brain, cortex and subcortex brain volume quantification from mice that received senolytic treatment (dasatinib (D) and quercetin (Q), DQ, n=6) or vehicle (n=8) for three months and compared to non-transgenic *Mapt^0/0^* mice (n=3). Data were analyzed with two-way ANOVA, Tukey’s *posthoc:* * *P* < 0.05; ** *P* < 0.005).

**Extended Data Figure 9.**
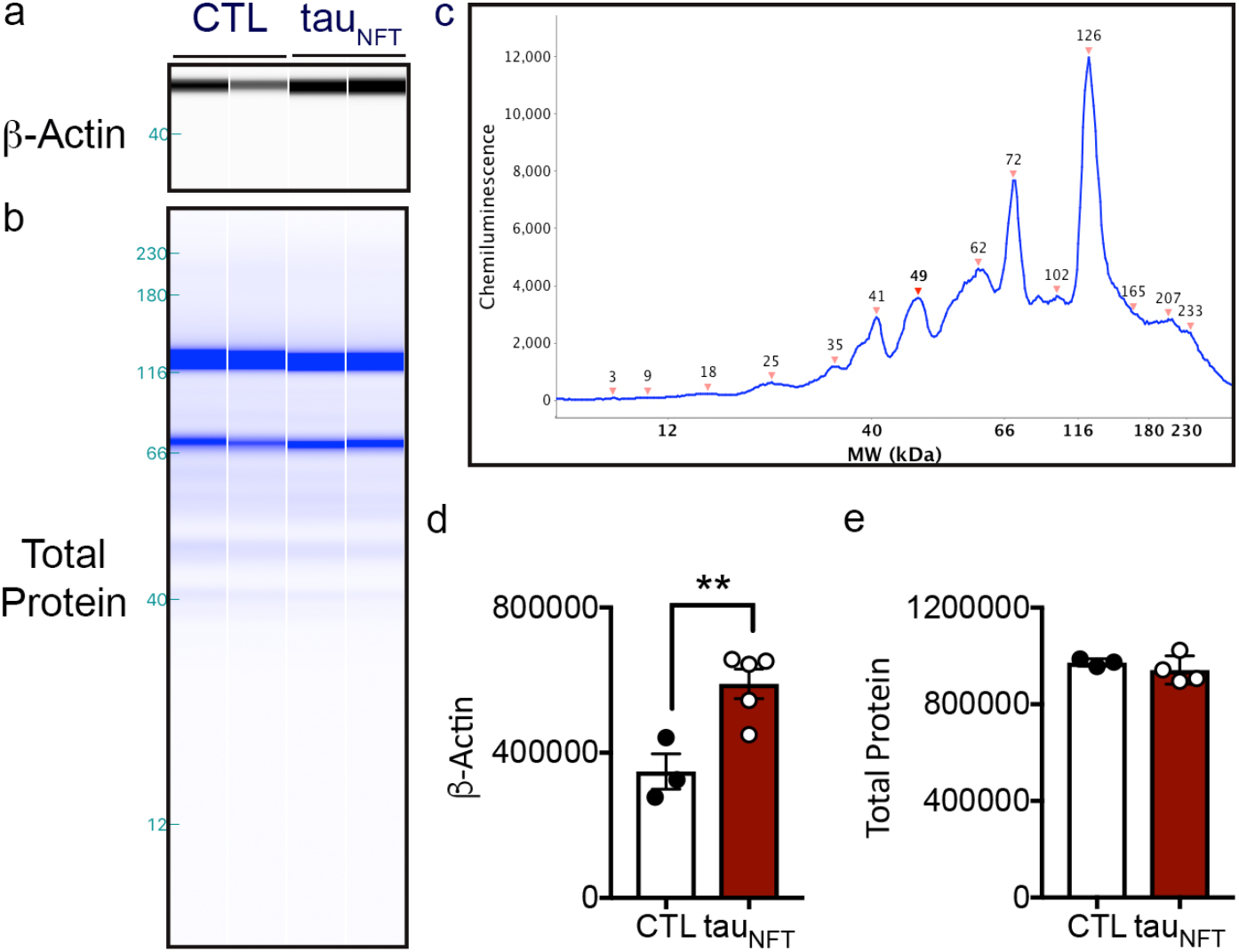
Total protein concentration was used as the internal loading control for protein quantification. (a) Representative capillary electrophoresis immunoblot using an anti-β-Actin antibody on brain homogenates from control or tau transgenic mice. (b) Representative capillary electrophoresis immunoblot used to detect total protein concentration. (d) Representative electropherogram generated from capillary electrophoresis total protein immunoblot. (d) Quantification of β-Actin and (d) total protein concentration. CTL: n = 3; tau_NFT_: n = 5; unpaired two-tailed t-test, β-Actin: ** *P* = 0.0097; Total Protein: *P* = 0.2916.

**Extended Data Table 1.** Human Postmortem Brain Characteristics.

|  | Control (n=10) | PSP (n=14) | p-value |
| --- | --- | --- | --- |
| Age at death (yrs) | 85.70 ± 2.81 | 83.86 ± 3.08 | 0.6765 |
| Sex (M/F) | 6/4 | 9/5 | N/A |
| Last MMSE score | 27.67 ± 0.87 (n=9) | 21.00 ± 2.02 | 0.0194 |
| Brain mass (g) | 1169 ± 17.83 | 1139 ± 43.21 | 0.5800 |
| Total Tangles | 4.03 ± 0.77 | 7.64 ± 0.66 | 0.0017 |

**Extended Data Table 2.** Primary Antibody Information.

| **Antibody** | **Supplier** | **Catalog Number** | **Clone Name** | **Lot Number** |
| --- | --- | --- | --- | --- |
| p65 | Cell Signaling | 8242 | D14E12 | 4 |
| phospho-Ser139 H2A.X | Cell Signaling | 9718 | 20E3 | 13 |
| HT7 | Pierce/Invitrogen | MN1000 | HT7 | QA208737 |
| NeuN | Millipore | MAB377 | A60 | LV1746157 |
| NeuN | Millipore | MAB377 | A60 | 2326372 |
| GFAP | Cell Signaling | 12389 | D1F4Q | 2 |
| PHF1 | Dr. Peter Davies | N/A | N/A | N/A |
| NeuN | Cell Signaling | 12943 | D3S31 | 1 |
| Histone 3 | Cell Signaling | 4499 | D1H2 | 9 |
| Synaptophysin | Cell Signaling | 5461S | D35E4 | 2 |
| Plp1 | Sigma | HPA004128 | N/A | N/A |

